# *F1000Research* TMATCH: A New Algorithm for Protein Alignments using amino-acid hydrophobicities

**DOI:** 10.1101/2019.12.16.878744

**Authors:** David Cavanaugh, Krishnan Chittur

**Affiliations:** Bench Electronics, Huntsville, AL 35805; Chemical Engineering Department, University of Alabama Huntsville, Huntsville, AL 35899

## Abstract

The identification of proteins of similar structure using sequence alignment is an important problem in bioinformatics. We decribe TMATCH, a basic dynamic programming alignment algorithm which can rapidly identify proteins of similar structure from a database. TMATCH was developed to utilize an optimal hydrophobicity metric for alignments traceable to fundamental properties of amino-acids. Standard alignment algorithms use affine gap penalties as contrasted with the TMATCH algorithm adaptation of local alignment score reinforcement of favorable diagonal paths (transitions) and punishment of unfavorable transitions paired with fixed gap opening penalties. The TMATCH algorithm is especially designed to take advantage of the extra information available within the hydrophobicity scale to detect homologies, as opposed to the probabilities derived from raw percent identities.

## Introduction

Protein and DNA/RNA sequence alignment algorithms are fundamental to modern bioinformatics. Sequence alignments are widely used in diverse applications such as phylogenetic analysis, database searches for related sequences to aid identification of unknown protein domain structures, classification of proteins and classification of protein domains. Additionally, alignment algorithms are integral to the location of related proteins in order to secure understanding of unknown protein functions, to suggest the folded structure of proteins of unknown structure from location of homologous proteins and/or by locating homologous domains of known 3D structure. [1] [2] [3] [4] Smith-Waterman (local alignment) and Needleman-Wunsch (global alignment) are exact match algorithms that use dynamic programming methods while BLAST and FASTA can be termed as “word” methods (e.g. local alignment of regions of high similarity) that while not guaranteed to find optimal alignment solutions are significantly more efficient and very useful in large-scale database searches. Dynamic programming alignment algorithms produce optimal/near optimal alignments from an optimization of the global alignment score derived from local alignment score optimization. The local alignment scores derive from a statistical transition probability matrix of nucleotides or amino-acids, depending upon the sequences being aligned.

We have developed an algorithm we call TMATCH that has adapted features of basic dynamic programming alignment algorithms. TMATCH was developed to utilize an optimal hydrophobicity metric for alignments as described elsewhere [5] Standard alignment algorithms use affine gap penalties. In TMATCH, the local alignment score reinforces favorable diagonal sequences that are paired with fixed gap opening penalties. The TMATCH algorithm is especially designed to take advantage of the extra information available within the hydrophobicity scale to detect homologies, as opposed to the probabilities derived from raw percent identities.

It is difficult to detect remote homologies in the so called twilight zone that have low percent sequence identities starting around 20-25 % and descending to around 10-15 %. The hydrophobicity scale we have described [5] is an excellent measure of sequence relatedness and identifies *similar* proteins much better than those derived using sequence identities, particularly in the twilight zone. A robust estimate of the hydrophobicity based sequence identity may be calculated directly from the global alignment score, which may be directly used in database searches, obviating the need to actually incur the overhead of maintaining the backtracking matrix (described later) and actually extracting the alignment. Low sequence identities, possessing statistically insignificant similarities by conventional measures, but having similar secondary structures can be identified using our algorithm, while not identified as statistically significant by other methods such as FASTA and Smith-Waterman.

### Approach

The TMATCH algorithm starts off with the dynamic programming algorithm used for alignments as is embodied in the Needleman-Wunsch global alignment algorithm. The TMATCH algorithm uses a fixed gap penalty and therefore abandons the notion of an affine gap penalty, which is problematical as there is no deep, underlying theoretical construct for choosing a specific affine gap penalty that derives from statistical theory and/or from protein function/structure. The TMATCH algorithm leverages the fact that local pair-wise sequences of high homology result in diagonal (upper left to lower right) traces in dot-matrix/dot-plot algorithms. When these local pairwise diagonal traces exist in the optimal/near-optimal alignment catchment basin, which is defined as an area about the dot-matrix major diagonal, they will contribute to and be included within the global/near global alignment traces within the dot-plot/dot-matrix. TMATCH captures these dot-plot/dot-matrix algorithmic properties by introducing the notion of score “rewards” for favorable (e.g. local alignment score optimization) cell-cell diagonal transitions and score “punishments” for unfavorable cell-cell diagonal transitions. Fixed gap penalties are assessed for horizontal or vertical cell-cell transitions. Pairwise comparisons of amino-acids in the alignment are done with an optimal hydrophobicity proclivity matrix developed in our companion hydrophobicity proclivity scale paper [5].

Methods are devised in this study to predict the percent hydrophobic match similarity of two protein sequences being aligned from the maximum score divided by the average number of pair-wise matches in the alignment. These methods allow a statistical relationship test to be performed without having to extract the aligned sequences and directly compute the percentage of hydrophobic match similarity. A statistical hypothesis test of homology is developed based upon percent hydrophobicity similarity match between two protein sequences using a Binomial fraction cumulative PDF. In order to speed computation and provide for maximum coverage of computational platforms and computational language implementations of the TMATCH algorithm, a hyperbolic function was developed to closely approximate the cumulative Binomial fraction PDF. Our computations of probability with the Binomial fraction PDF and hyperbolic approximation function calculate tail areas at and above a threshold Zb, as opposed to calculating an area less than or equal to the threshold Zb.

## 1 Methods

We describe the TMATCH algorithm using two, six residue synthetic poly-peptide sequences using amino acid hydrophobicity proclivity indices we have created [5]. These sequences are VLSEGE (sequence 1, shown as a row) and VHLTPE (sequence 2, in the column)

The calculations start with the creation of a match matrix and this is shown as Table 1. We then create a score matrix (Table 3) and a back track matrix (Table 4) These three matrices are used to create the final alignment and a score.

**Table 1.**
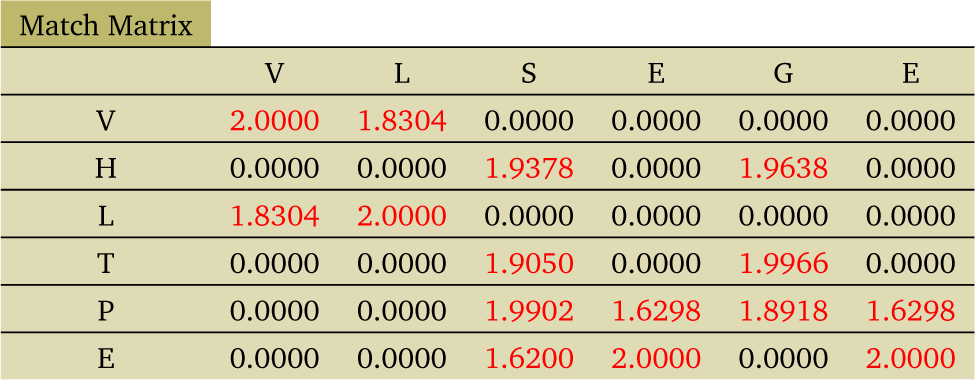
Match Matrix

Each matrix (as seen in the Tables 1, 3 and 4) organized around the row (search protein) string and each of the incoming (match protein) column strings being aligned against the row (search) string. Each matrix column corresponds to one character of the row string and each row character corresponds to a character of the column string. The match matrix (table 1) has scaled delta hydrophobic proclivities which represent similarities between the amino-acids being compared, where a score of 2 is a perfect match and a score of 0 is declared a non-match. Scaled delta hydrophobic proclivities below a threshold are windowed by forcing cell values to zero for each scaled delta hydrophobic proclivity less than the threshold value. The match matrix will contain diagonal sequences representing contiguous sequences of amino-acids with high similarity, comparable to what a dot plot will do.

The score matrix (Table 3) is derived like the recursion relations of the classic dynamic programming algorithm, such as with the Needleman-Wunch algorithm. Concomitantly with generating the score matrix, the backtracking matrix (table 4) is generated with each cell corresponding to the same cell of the score matrix, and each backtrack matrix cell value indicating which was the previous cell path chosen in the score matrix (from left, up or upwards left diagonal). The backtracking matrix cell corresponding to the maximum value cell in the score matrix is then used to extract the alignment, moving from the string right (end) and growing towards the string left (front). The direction taken in the backtracking matrix will determine the growth of the aligned row/column strings either by taking amino-acid characters from the row or column strings respectively, or by introducing gap characters as necessary.

The match matrix match values are calculated by first calculating the difference in hydrophobicities between the amino-acids in each row and column intersection, subtracting the absolute value of this difference from 1.0, and if the difference is greater than a threshold value, is multiplied by a weighting factor. We chose the inverted hydrophobicity proclivity distance (1 minus the hydrophobicity proclivity distance) threshold value of 0.8 (hydrophobicity proclivity distance of 0.2) and a weighting factor of 2.0, for a match value distance threshold value of 1.6. Therefore, the match value threshold of 1.6 comes from a maximum allowed fuzzy match hydrophobicity proclivity distance of 0.2. The Mw (weighting factor) of 2.0 derives from an initial, discrete version of the TMATCH algorithm designed to work with plain alphanumeric string comparisons. The values of the gap penalty, the transition reward and the transition penalty were also empirically and iteratively determined in order to yield optimal alignments under different and taxing conditions. Details of this portion of the work have been necessarily omitted from this current paper for brevity sake. The portion of the work we are now showing serves to illustrate the reasonableness of the fitted reward/penalty values. The result of these calculations are shown in Table 1.

Here are some sample calculations using the hydrophobic proclivity values from the table 2 as seen in [5].

**Table 2.** Table of Regression Fitted Hydrophobic Proclivities

| Residue Amino Acid | Hydrophobicity (H) | Regression Fitted $\Delta G$ |
| --- | --- | --- |
| F (Phenylalanine) | 0.0688 | 2.5658 |
| L (Leucine) | 0.0579 | 2.6095 |
| I (Isoleucine) | 0.0349 | 2.7022 |
| M (Methionine) | 0.2213 | 1.9528 |
| V (Valine) | 0.1427 | 2.2687 |
| P (Proline) | 0.7123 | -0.0212 |
| T (Threonine) | 0.6599 | 0.1895 |
| S (Serine) | 0.7074 | -0.0018 |
| A (Alanine) | 0.4925 | 0.8624 |
| Y (Tyrosine) | 0.4523 | 1.0237 |
| H (Histidine) | 0.6763 | 0.1232 |
| Q (Glutamine) | 0.8692 | -0.6522 |
| N (AsparagiNe) | 0.8350 | -0.5148 |
| K (Lysine) | 0.9651 | -1.0376 |
| D (Aspartic Acid) | 0.9157 | -0.8393 |
| E (Glutamic Acid) | 0.8974 | -0.7657 |
| C (Cysteine) | 0.2650 | 1.7769 |
| W (Tryptophan) | 0.3403 | 1.4742 |
| R (ARginine) | 0.9091 | -0.8126 |
| G (Glycine) | 0.6582 | 0.1961 |

We begin with the calculations for PS (row value of 5, column value of 3)

1. Calculate1.0 − *abs*(hydrophobicity P − hydrophobicity S) which is (using values from table 2) 1.0 *− abs*(0.7123 *−* 0.7074) which is 0.995
2. Since 0.995 is greater than 0.8, match value is 2.0*0.995 which is 1.99 shown in Table 1

We next show the calculations for LS (row 3, column 3)

1. Calculate 1.0*−abs*(0.0579*−*0.7074) which is 0.3505
2. Since 0.3505 is smaller than 0.8, the match value is zero.

The entire Table 1 is calculated this way. Owing to the nature of the TMATCH algorithm, there is a trough (catchment basin) of high likelihood constraining possible alignment paths, running from the top left to the bottom right corners. Horizontal or vertical blocks (e.g. with a non-zero match value, shown in red) mostly surrounded by white space (e.g. 0 match value) generally represent regions of gap insertions, should the alignment path run through those blocks. The “isolated” vertical/horizontal red blocks represent amino-acid regions of high similarity, which will likely be part of the alignment path, if they can be diagonally connected on one side or the other and lie within the alignment catchment basin. The alignment path (corresponding to gaps) may pass through white (zero value) cells, within the catchment basin, between islands of red (non-zero value) cells.

Our algorithm assigns a fixed gap penalty of 0.8. Each cell in a dynamic programming score matrix reflects the cumulative score from the start to that point, as the dynamic programming algorithm finds the best path through the matrix by passing through all of the best (or near optimal) local scores along the catchment basin.

A cumulative score matrix is calculated next (shown as Table 3)

**Table 3.**
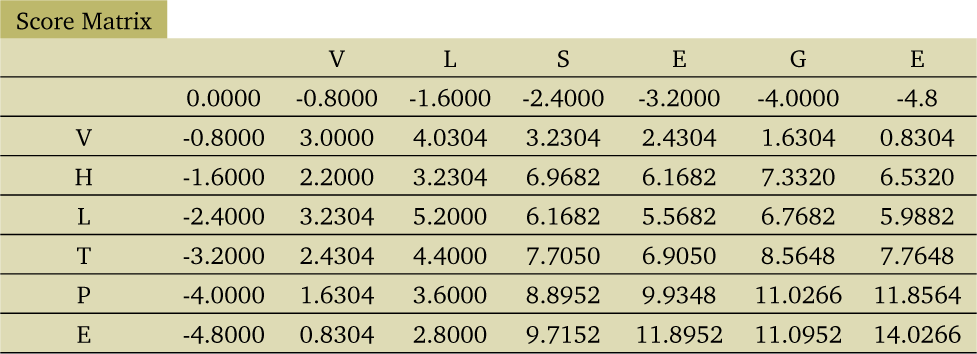
Score Matrix

The calculation starts at the first row and first column (in this case at V & V). The score value at each cell is calculated using information from the previous diagonal cell and those from the cell in the previous row (up) and the cell from the previous column (left). At each step of the calculation, the current cell in the score matrix is the same cell being accessed in the match value matrix. The starting score value in the score matrix is set to zero. Subsequently, the score assigned to each cell of the score matrix comes from the highest score achieved by selecting an immediately previous cell on the left hand side, on the upper left hand diagonal side or the on the upper side. For each score matrix cell processed, the first calculation step adds the match matrix cell value corresponding to the current score matrix cell to each of the 3 possible previous score matrix cells (left, diagonal or up), each of which could be one of the three putative paths into the current score matrix cell. In step two of each current score matrix cell calculation for the left and up cell paths, a fixed gap opening penalty of −0.8 is added. The second step of the current score matrix cell calculation for the diagonal cell path is a more complex in that the potential transition is assessed a penalty (−1.4) for a bad transition (match value matrix diagonal cell value *>* the current match value matrix cell value) or granted a reward (+1.0) for a favorable transition (match value matrix diagonal cell value *<*= the current match value matrix cell value). The second step of the current score matrix cell calculation for the diagonal cell path reward/penalty is added to the diagonal path calculation. The path chosen is that resulting in the highest score for the current score matrix cell. The first row only has left hand entry transitions and the first column only has entry transitions from the up direction, with concomitant application of the gap penalty (−0.8) for each cell transition. Effectively, the calculation proceeds in an iterative fashion matching that of the traditional dynamic programming algorithm.

To illustrate these algorithmic steps we calculate the upper left 6 cells of the score matrix, with a couple of examples of where the back track matrix trace will go through and how they are determined. Each cell address is reported as (column label, row label), such that the third column & second row is cell LV:

1. (The initial entry into the score matrix is 0)
2. VV: 0 (upper left initialization) +1 (favorable diagonal transition) +2 (equiv match matrix cell) =3
3. VH: 3 (upper cell) −0.8 (gap penalty) =2.200
4. LV: 3 (cell VV) +1.8304 (equiv match matrix cell) - 0.8 (gap penalty) =4.0304
5. LH: (from left): 2.2000 (cell VH) +0.0 (equiv match matrix cell) −0.8 (gap penalty) =1.4000
6. LH (from up): 4.0304 (cell LV) +0.0 (equiv match matrix cell) −0.8 (gap penalty) = 3.2304
7. LH (upper-left): 3.000 (cell VV) +0.0 (equiv match matrix cell) −1.4 (bad diagonal punishment: 2>0 in match matrix) =1.6000
8. LH: 3.2304 (highest score from up, equiv back track cell value =1)
9. VL: 2.2000 (upper cell) +1.8304 (equiv match matrix cell) −0.8 (gap penalty) =3.2304
10. LL (from left): 3.2304 (cell VL) +2.0000 (equiv match matrix cell) −0.8 (gap penalty) =4.4304
11. LL (from up): 3.2304 (cell LH) +2.0000 (equiv match matrix cell) −0.8 (gap penalty) =4.4304
12. LL (upper-left): 2.2000 (cell VH) + 2.0000 (equiv match matrix cell) +1.0 (favorable diag reward, 0<2 in match matrix) =5.2000
13. LL: 5.2000 (highest score from upper-left, equiv backtrack matrix cell value =0)

The successive building of cumulative score matrix cell values in Table 3 are then used to build the alignment in the backtrack matrix Table 4 from the score matrix local cell maximum computed scores, where there is a direct location correspondence between each cell in the score matrix and backtrack matrix. For each cell in the score matrix and its backtrack matrix analog cell, the algorithm for building the backtrack matrix will mirror the score matrix cell entry path from the direction resulting in the maximum value for the current score matrix cell. If the current maximum score matrix cell value was arrived at from the previous diagonal, a value of 0 is assigned to the corresponding backtrack matrix cell. If the current score matrix cell value was arrived from the score matrix cell on the left, the backtrack matrix cell value assigned is −1 (meaning there will be a gap in the column and the row symbol taken), and if the score matrix cell value was arrived at from the upper cell, the value of +1 is assigned to the backtrack matrix cell (gap in the row and the column symbol is taken).

**Table 4.** BackTrack Matrix

| BackTrack Matrix |  |  |  |  |  |  |
| --- | --- | --- | --- | --- | --- | --- |
|  | V | L | S | E | G | E |
| V | 0.0000 | -1.0000 | -1.0000 | -1.0000 | -1.0000 | -1.0000 |
| H | 1.0000 | 1.0000 | 0.0000 | -1.0000 | -1.0000 | -1.0000 |
| L | 1.0000 | 0.0000 | 1.0000 | 0.0000 | 0.0000 | -1.0000 |
| T | 1.0000 | 1.0000 | 0.0000 | -1.0000 | 0.0000 | -1.0000 |
| P | 1.0000 | 1.0000 | 1.0000 | 0.0000 | -1.0000 | -1.0000 |
| E | 1.0000 | 1.0000 | 1.0000 | 0.0000 | -1.0000 | 0.0000 |

V-LSEGE Aligned row string
VHLTP-E Aligned column string

The alignment itself is extracted by starting with the backtrack matrix cell corresponding to the score matrix cell that had the maximum cell value and backtracking to each of the next (previous) cells as indicated by the stored path direction in each backtrack matrix cell. The alignment path follows backward though each cell obtained by the successive recorded entry directions. Concomitantly, the row and column protein alignment strings grow from right to left as the alignment is extracted from the back-track matrix. For a 0 backtrack matrix cell value, both the next (from the right hand side or end position) row and column protein string characters are taken into the growing (right to left, end to front) aligned row/column strings. For a −1 backtrack matrix cell value, a gap (−) character is put into the front (left) of the growing aligned column string and the next (end) character from the row protein string is taken for the front (left) of the aligned growing row string. For a +1 backtrack matrix cell value, a gap (−) character is put into the front (left) of the growing aligned row string and the next (end) character from the column protein string is taken for the front (left) of the aligned growing column character string.

The red highlighted cells in table 4 above shows the back-track trace used to recover the gapped alignment strings, shown at the bottom of table 4. Under each alignment string corresponding to each amino-acid is a match value as represented in table B above. A match value distance of two represents a perfect match. In this case, the total of the match values is 14.0266. While indicative of how the two sequences are similar, this sum match value is not as useful as the score matrix cell values in database searches, or looking for proteins with similar properties.

In the interest of large database searches it is desirable to minimize the computational load to keep the database searches within reasonable time frames. We show how this can be done by calculating a set of metrics from the alignment.

Now we turn to the rationale and description of the statistical significance test of the aligned protein character strings. We have used the serine protease protein family data from Pearson [4] to calibrate the binomial distribution (our selected distribution) expected parameter P (derived below) for amino acid pair matches from random search and match sequences being aligned. The statistical hypothesis test null hypothes used is that the two aligned sequences are not related (they are random), so the post alignment %H matches are less than or equal to the fitted P (parameter) value, which will result in the the binomial distribution function right hand tail area of greater than 10% (i.e. Bn(Z) > 0.1), resulting in the declaration that two random sequences are unrelated.

The two sequences chosen to bracket Bn(Z), which is approximately = F(Zb’) = alpha = 0.1 (derived below) are in the region where the Smith Waterman statistical significance function E() values transition into statistical non-signficance. The significance function E() used for the Smith-Waterman alignment is considered to be marginally significant around 0.0001 to 0.01, whereas the same region for the binomial Bn(Z) PDF is around 0.10. As can be seen from table 5 below, the Bn(CO2_HUMAN) and Bn(CFAB_MOUSE) values actually bracket the alpha =0.10 value for F(Zb’) corresponding to the E() transition threshold of statistically insignificant alignment. The TRYP Bovine Serine Protease was used as the search protein [6] [4]. Table 5 below also shows the percent identity value to be approximately the square root of the hydrophobicity fuzzy match percent (i.e. there is more information in the hydrophobicity index).

**Table 5.** Match

| Protein | %H match | % identity | $F(Zb')$ | S.W. $E()$ |
| --- | --- | --- | --- | --- |
| CO2_HUMAN (P066811) | 39.84 | 17.5 | 0.1319 | 1.00E-04 |
| CFAB_MOUSE (P041851) | 40.55 | 21.25 | 0.0960 | 1.00E-02 |

Since the hydrophobicity proclivities are variables (e.g. not discrete), in order to make logic comparisons it is necessary to stipulate an interval definition of similarity for a "fuzzy match" indicating a pairwise match owing to similarity between two amino acids being compared. Two definitions of similarity were chosen for loose and strict fuzzy matches, H ± 0.2 and H ± 0.067 respectively. The fH values reported in table 5 are the average of the strict and loose fH values per this definition, which is used for the calculation of the TMATCH F(Zb’) significance function values. A compromise bin width of 0.2 was used for making a theoretical prediction of the cumulative Binomial PDF expected/average percent. As mentioned above, inspection of the distribution of the hydrophobicity proclivity values shows that that this bin width yields approximately 4 amino-acids per bin, or the whole 20 amino-acids over the whole interval of 0 to 1, hence the choice of 0.2 for the bin width.

The null hypothesis is that a search sequence is not statistically related to the match sequence. We use the bionomial probability distribution function (PDF) in constructing a test for the null hypothesis. Examining the hydrophobic proclivity values, we can see that the 20 amino acids are distributed almost evenly over the whole interval from 0 to 1, 4 amino acids in each of the 5 interval bins of width 0.2. Using this observation, we start with a search sequence as given and a putative match sequence alignment is assumed to result in random (uniform distribution) pairwise amino acid match ups, where two amino acids being compared from search and match sequences are declared to be matching if both amino acids being compared fall into the same interval of the 5 predefined intervals. The bin counts after the alignment of random search and random match sequences should result in the 5 interval bin counts having nearly the same values. This result would then suggest that 5 equal intervals used to determine a pairwise amino-acid match would result in a Binomial PDF P parameter of around 0.2 for the statistical determination of “non-relatedness” from the alignment of two random amino-acid sequences.

There are two ways to construct a detailed probability argument for calculating the theoretically expected Binomial average and assemble the null hypothesis. Firstly, the search sequence is a *known* or given and each subsequent sequence to which it is compared using an alignment can be considered for the sake of this analysis to be a random (uniform distribution) sequence of amino-acids. Furthermore, we can make the assumption that the statistical paradigm we are using is sampling with replacement, so the idealized probabilities can be used. Therefore, for any amino-acid selected from the search sequence is considered to be fixed for the comparison purpose and is located in one of the fixed width bins. The amino-acid to which it is compared from the comparison sequence is either in the same bin or not. If the selected comparison sequence amino-acid falls into the same bin, it is treated as a fuzzy match. In actuality, the fuzzy match bin is centered about the hydrophobicity proclivity of the selected search sequence amino-acid, but for the sake of simplifying the calculations five fixed bins have been assigned. For the purposes of selecting the expected average of the cumulative Binomial PDF, the null hypothesis chosen here is that the selected search sequence amino-acid is fixed, non-random and that the comparison or match sequence amino-acid to which it is compared is random and a match occurs when the selected comparison/match sequence amino-acid randomly drops into the same bin as the fixed search sequence. Under this scenario, there are 4 ways out of 20 (20% chance) that a fuzzy match occurs as there are 4 possible randomly selected amino-acids in the comparison/match sequence which could fall into the same bin as the fixed search sequence (e.g. sampling with replacement).

A second calculation approach may be made by assuming that both the selected search sequence amino-acid and the matched amino-acid in the search string are both random and each amino-acid would independently and randomly have dropped into the same bin. The probability of the two amino-acids randomly dropping into the same bin are (4/20)*(4/20)=4%, but there are 5 ways that this could happen, so the actual probability (average) over all comparisons would be derived from the fact that there are 5 ways (5 bins) that the random match could occur yielding a N*P=5*4%=20%. Both calculations yield the same random match probability for any amino-acid pair comparison between the aligned search sequence and aligned comparison/match sequence. The percentage match under random conditions between the two sequences would then simply be the expected probability value on a amino-acid pair comparison, neglecting the contribution of insertion characters which will affect the actual results seen in practice. However, for the purposes of calculating the expected Binomial PDF average of 20%, some reasonable simplifying assumptions have been made and we see from fitting the cumulative Binomial PDF average from the Pearson [4] Serine Protease data set that the 18.6% is close to the expected 20% as derived above.

As discussed above, the actual P (parameter) value used for the binomial distribution (Bn(Z)) (Z score average percent to standardize Bn(z)) was calibrated from a theoretical initial estimate of 0.2 with a subsequent iterative fit using a Tryptophan like Serine protease data set known to be significantly related by a Smith-Waterman alignment in Pearson [4]. This Tryptophan like Serine Protease family from Pearson [4] was used to calibrate/fit the cumulative binomia PDF Bn(Z) random amino-acid match parameter P on the interval 0.15 to 0.2 using the Bn(Z) interval bracketed with the CO2_Human and CFAB_Mouse Serine Proteases to yield F(Zb’) values approximating the marginally significant ("probably related") to non-significant transition alpha value of 0.10 from the F(Zb’) PDF. The F(Zb’) interval of alpha = 0.10 is bracketed by f(CO2_HUMAN; P06681-1) = 0.1319 and f(CFAB_MOUSE; P04186-1) = 0.0960. Similarly, we see in table 5 that these two Serine protease enzymes also span the E() transition from marginally significant to non-significance. These F(Zb’) values have been reported in a table in the companion application paper [6] where we have applied the TMATCH algorithm to the set of Tryptophan like Serine Proteases characterized by Pearson [4]. The Bn(Z) function is fitted with a 5 parameter hyperbolic significance function F(Zb’) (see below). Bn(Zb) or F(Zb’) values above 0.1 are at the threshold value where we say that two aligned proteins are not statistically related. The average P (parameter) value is the assumed maximum percent threshold at which two random sequences should be deemed to be unrelated as the significance function F(Zb’) (or Bn(Zb)) will result in a value above the alpha value of 0.1.

The hyperbolic significance function calibration results we report above, was essentially replicated with datasets representing multiple protein families: DNA Polymerase B enzymes, G proteins, Glutathione proteins and Rhodopisin/GPCR proteins [6].

We present the following definitions:

1. fHe =0.186 (fraction H match expected for random matches Calibration algorithm is described above)
2. fHs is the loose fuzzy hydrophobicity match between amino-acid pairs being compared, which is calculated as the fraction of matches where the absolute value of the difference in hydrophobicity proclivities is less than or equal to 0.2

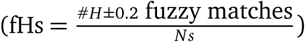
3. fHr is the strict fuzzy hydrophobicity match, which is calculated similarly to fHs, but here the absolute value of the difference in hydrophobicity proclivities is less than or equal to 0.067. 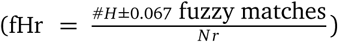 and,
4. Ns is the number of characters in the aligned row/column strings and Nr is the number of characters in the row/search string (and Ns *>*= Nr).

Experimenting with a number of sequences, we concluded that the fHs statistic is too sensitive to significant mismatches in sequence length and the average of fHs and fHr gave a more stable yet still sensitive statistic that reflects real protein primary sequence matches/mismatches.

We next calculate a Z score transformation of the form: (value-average)/standard deviation. The statistics Zb and Zb’ defined below are used as part of the calculation of a hyperbolic statistical significance function F(Zb’) fitted to the cumulative binomial probability density function Bn(Zb,fHe), serving for the TMATCH sequence alignment statistical significance calculation:

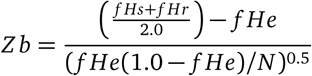

Where N is the number of amino-acid pair comparisons

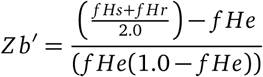

Where fHe was calibrated to be 0.186 from simulations with the protein kinase data set from [3] and F(Zb’) is then calculated as:

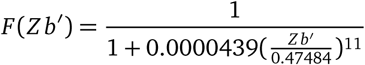

Which is a hyperbolic function that very closely approximates a binomial distribution Bn(Zb, 0.186) and used to calculate the significance of the aligned row/column string matches. An average score Sa is calculated from the alignment:

Sa = (maximum alignment score)/(Mw x median protein length)

Where the median protein length is simply the average length of the two proteins being aligned with respect to each other and the weight Mw is 2.0. Our simulations with the Serine protease set show that a plot of the Sa^1.56^ (which we define as fHst) versus fHs is a very good straight line. Thus, it is possible to relate an average score obtained during computations to calculate fHst an estimate of the fHs metric for database searches. The fHst metric may be substituted for the average of the fHs and fHr metrics in the Zb calculation above for use during the first pass search for suitable match candidates to the search sequence protein.

Below is the statistical significance (alpha) interpretation of the F(Zb’) hyperbolic significance function:

1. F(Zb’) *<*= 0.1, probably related
2. F(Zb’) *<*= 0.05, related
3. F(Zb’) *<*= 0.01, strongly related
4. F(Zb’) *<*= 0.001, very strongly related

The fHst is a very good fit of fHs for fHs values between 20% and 70%. For fHs values above 0.85 in some data sets, the fHst calculation fHst = Sa^1.56^ can diverge enough to result in pathological behavior of fHst, where calculated fHst values may exceed 100%, because the average alignment score Sa is greater than 1. For data sets leading to pathological fHst behavior (Sa *>* 1), two alternate equations for fHst may be used to represent a better fit of the data across the whole range of possible fHs values

1. fHst = Sa^1.495^ where Sa *<*= 0.69
2. fHst = Sa^1.495^/1.151 where Sa *>* 0.69

The average score Sa can be used to estimate the %H match based upon the fHst^0.5^ statistic (Figure 2). The fHst statistic is adequate for initial database searches and multiple alignments, as it is fairly conservative statistic. Searching for related sequences in the protein databases would be done using sequence fuzzy matching using fHst and the resulting statistical scoring would be calculated from the F(Zb’) based significance function. A search sequence with fHst fuzzy matches on the interval of 5%-10% can uncover distantly related, but homologous protein sequences of percent identities in the twilight zone range of 22%-32% identity. Even modest improvements in algorithmic efficiencies make large differences when searching many thousands of records. Efficiencies of scale derive from the ability to drop the backtrack matrix calculation and alignment extraction step, thereby making a significant time difference with TMATCH for database searches, as the average alignment score Sa can be directly used to calculate fHst and F(Zb’).

## 2 Discussion

In Figure 2, we show the relationship between the average of fHs^2^ and fHr^2^ and the percent identity (the fraction of exactly matching aligned amino-acids) from the set of serine proteases. This relationship cannot be an artifact of the TMATCH algorithm, but represents something about the underlying relationship between homologous proteins. In the limit of homology (e.g. the twilight zone 25% identity), the essential core of amino-acids needed for a particular function is revealed within the homologous amino-acid positions within the aligned sequences. For example, two proteins of 25% identity (as seen in Figure 2) in reality reflect a similarity of 50%. In this way we see that the apparent dilemma of divergent proteins (based upon the primary sequence) sharing essentially the same fold and function is in fact not a problem at all, because the biological function and structure is determined by a strongly homologous minority of amino-acids and supported by a large remaining fraction of similar amino-acids at analogous positions within the homologous proteins.

Presumably, the essential core of amino-acids represents in some way the minimum description of the tertiary structure of a given family of homologous proteins. The amino-acid hydrophobicity represents a similarity scale reflecting the actual physico-chemical similarities between amino-acids, thereby reflecting the maximum deviation allowable in amino-acid substitutions. The definition of the fHs statistic provides for reflection of information about the number of gaps introduced into an alignment. The definition of the fHr statistic reflects information regarding a more strict comparison between the search sequence and another protein.

As proteins in a homologous protein diverge within a given protein family, simultaneously new amino-acids are being introduced as existing amino-acids are being substituted with similar amino-acids of comparable sizes, thereby on average preserving the replaced amino-acid aggregate volumes in order to preserve efficient packing within the folded core of globular proteins. Using the strict definition of hydrophobic similarity range of fHr along with an inspection of table 2, would suggest that any given amino-acid could be substituted on average by 3-6 other amino-acids.

If the average score Sa is used to estimate the %H match based upon the fHst statistic, then fHst would be estimated as described by the equations above. The fHst statistic is adequate for initial database searches and multiple alignments, as it is fairly conservative statistic. Searching for related sequences in the protein databases using sequence fuzzy matching based statistical scoring calculated from the F(Zb’) based significance function on the interval of 5%-10% will likely uncover distantly related, but homologous protein sequences. Even modest improvements in algorithmic efficiencies make large differences when searching many thousands of records. Efficiencies of scale derive from the ability to drop the alignment extraction step thereby making a significant time difference with TMATCH for database searches, as the average score can be used to calculate fHst.

In discussing the three figures in this paper it is important to note that calculated statistics will be discussed as percentages, which should be considered identical with their decimal fraction equivalents that has been used indicated using a leading ‘f’ for most variable names defined within this paper.

Figure 1 shows the modified fHst according to the defining equations above with a threshold value of 0.69 for the average score Sa. The linearity with a 469 member DNA polymerase B dataset (see the companion applications paper [6]) is excellent, with the regression line possessing an obvious 45 degree angle and passing through zero. The modified % Hst calculation eliminated fHst values above 100 % resulting from Sa values above 1. The excellent linearity between Hs (calculated from the extracted, aligned strings) and the fHst statistics (calculated from a power law relation with the average maximum score) shows the fundamental consistency of the alignment algorithm and chosen weights with the underlying amino-acid proclivity metric and the definition of fH strict and loose fuzzy matches. Figure 1 illustrates the justification for substituting the fHst statistic in the Zb equation in place of the average of the fHs and fHr statistics, thereby calculating the F(Zb’) test statistic for the relationship between two proteins directly from the score matrix (using Sa) without incurring the overhead of the alignment extraction.

**Figure 1.**
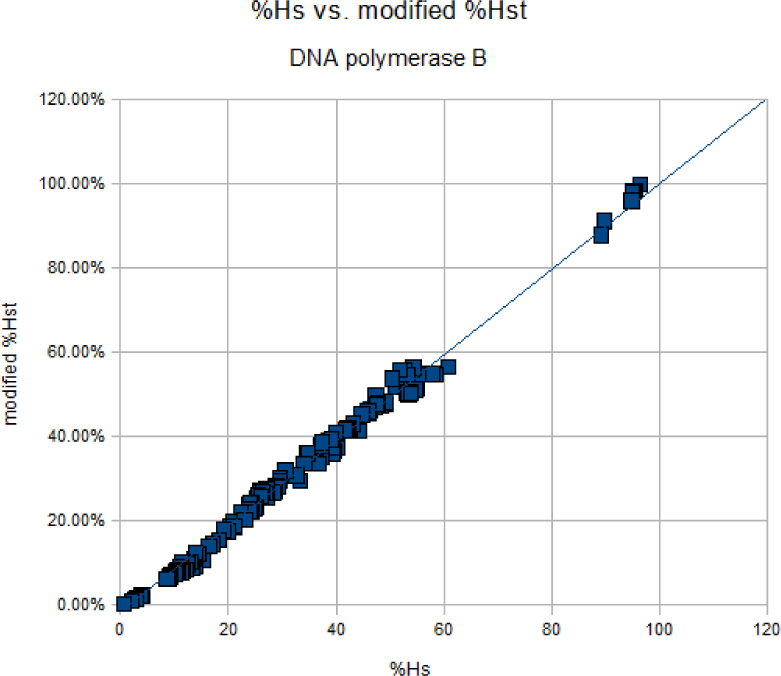
Fit of %Hs to %Hst. (R=0.9922, R^2^=98.44%)

As may be observed from figure 2, there is a strong relationship between the % identity (e.g. the fraction of exactly matching aligned amino-acids) and the % hydrophobicity match defined with a definition, such as fHs or fHr. Averaging the square of the fHs and fHr metrics results in a strongly linear (R^2^ =82.2 %) relationship with the % identity. We can see that this relationship is more than an artifact of the TMATCH algorithm itself and represents essential components of the underlying relationship between homologous proteins. Mathematically this result can be explained as the number of row string matches and the number of column string matches serving as linear Manhattan distances corresponding to two legs of a right triangle, and the percent identity serving analogously to the square of the hypotenuse of the right triangle. In this specific case, the averaging fHs and fHr is analogous to setting the legs of the right triangle to be equal.

**Figure 2.**
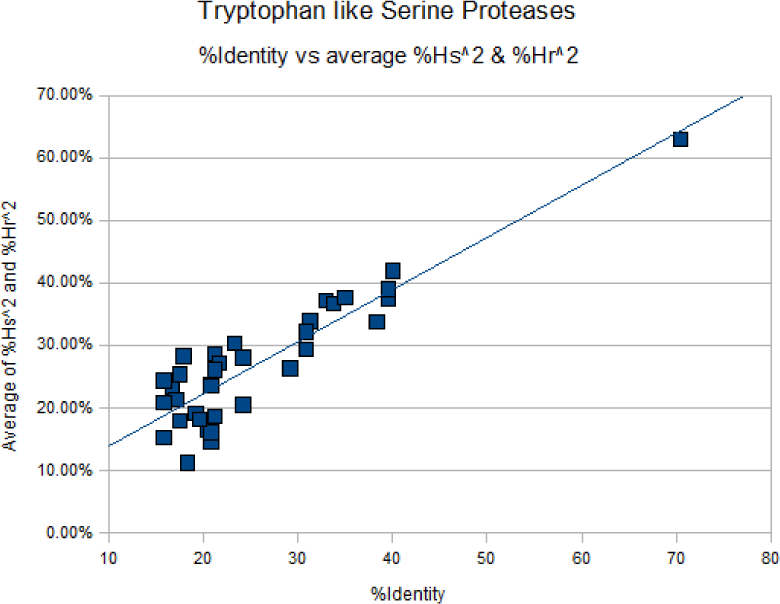
% Identity estimated from the average of fHs^2^ and fHr^2^ Bovine protein search sequence. (R=0.9068, R^2^=82.24 %) Sequences from [4]

Not all programming languages will have a convenient library function to implement the cumulative binomial function. Also, due to the nature of statistical cumulative PDF functions, they will take significant amounts of computation time if used hundreds or thousands of times as in large protein sequence database searches. Therefore it is desirable to estimate the cumulative binomial distribution Bn(Zb) in the form of a fitted function F(Zb’), using the form of a general five parameter hyperbolic function as stipulated above.

The plot in figure 3 is showing the relationship between the percent hydrophobic fuzzy match and the probability of two sequences not being related using one minus cumulative PDF functions, to invert the cumulative PDF’s to give the correct sense of the tail. We see that 1-Bn(35%, Zb) versus the normal approximation 1-N(Zb) shows an excellent fit. We also see that one minus the cumulative binomial function (Zb’,35%) with the fitted five parameter hyperbolic function F(Zb’,18.6%) is a very good fit up to 20%, which is really the whole range that the hyperbolic approximation is designed to fit with F(Zb’,18.6%), but also notice that the F(Zb’,18,6%) rises faster than the inverted cumulative binomial PDF. The centering/location of the two curve knees and the angle of each function roll off matches to very high degree as can be seen in figure 3. For the purposes of figure 3, an N of 100 was used. Substituting other N’s in the range of typical globular protein sizes shows that the cumulative binomial function to hyperbolic function relation is still reasonably close and should not affect the validity of the approximation.

**Figure 3.**
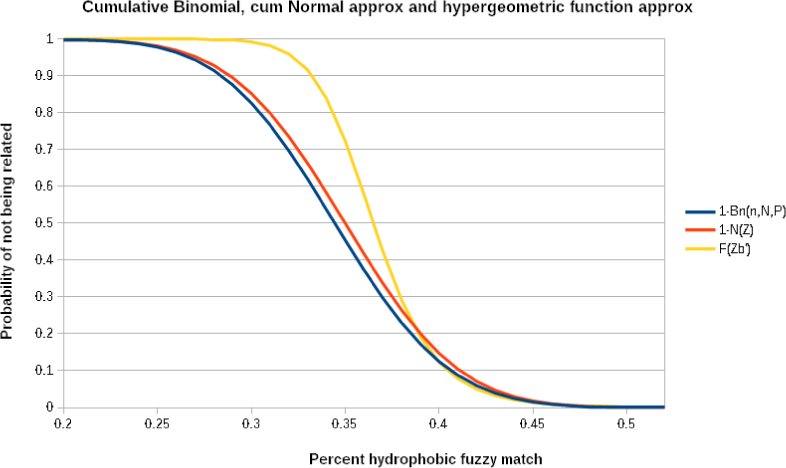
(%Match, 1-N(Zb)) is the red curve, (%Match, 1-Bn(35%) is the dark blue curve and (%Match, F(Zb’) is the yellow curve)

The fitting of the five parameter F(Zb’) function to the Bn(Zb) function was done manually and iteratively to minimize the sum of the square of errors using a number of select basis points on the two curves. The shape of the cumulative Binomial function Bn(Zb) function in figure 3 is actually determined by computing the outer single tail area past the specific Zb or Zb’ values; therefore this presentation of the cumulative PDF is an inverse of the typical presentation formed by computing the cumulative area less than the specific Zb (or Zb’) value, so the sum of these two presentations is equal to 1 for all Zb >= 0. The right half of the Bn(Zb) curve area has been divided by 0.5 to normalize the entire right half area to one so as to effectively represent the entire cumulative PDF by artificially precluding any Zb value from being less than zero.

The F(Zb’,18.6%) was calibrated against the E() expectation function used in BLAST and in the tryptophan like serine proteases and alignments reported in [4]. The Zb statistic was used in the cumulative binomial (and normal approximation) and it represents a traditional Z score statistics transformation. For the cumulative binomial function a value of Zb corresponding to 35% aligns the F(Zb’) and 1-Bn(Zb) significance functions. The Zb’ transform denominator is the variance of a population average 18.6%, rather than the traditional standard deviation, which was a necessary modification to get the hyperbolic significance approximation function slope to match that of the one minus cumulative binomial function and that of the E() significance function. Note that 0.35%^1.6022^ =18.6%, which is near a theoretically expected exponent of 2 reflecting a variance versus standard deviation relationship.

## 3 Conclusion

The essential core of amino-acids needed for a particular function is revealed within the homologous amino-acid positions within the alignment sequences. In the limit of homology (e.g. the twilight zone), it can be seen that similar structures have been achieved in nature by substituting similar amino-acids, especially for the essential core set of amino-acids. We have shown in the companion hydrophobicity paper [5] as well as in the results of this paper, that our hydrophobicity proclivity scale provides an excellent numerical definition of similarity between two amino-acids. The TMATCH algorithm mixes the key algorithmic features of dynamic programming algorithms and the key aspects of the dot-matrix/dot plot method in a way to take excellent advantage of the extra structural/functional information implicit within our hydrophobicity proclivity scale. The current form of the TMATCH algorithm described in this paper has been optimized for efficient search of large protein databases returning match results of higher sensitivity and reliability in cases of low sequence percent identity than searches based upon finding sequences of local homology because the TMATCH algorithm can access information about the overall structural information of the protein sequences being compared since TMATCH is a global alignment algorithm in the same vein as is the Needleman-Wunch algorithm.

